# Functional aberration of cortical neuronal network induced by Aβ42 oligomer

**DOI:** 10.1101/2023.03.13.532496

**Authors:** Dulguun Ganbat, Jae Kyong Jeon, Sang Seong Kim

**Author notes:** Correspondence to: Sang Seong Kim, Hanyang University, Gyeonggi-do, Republic of Korea.

## Abstract

Alzheimer’s disease (AD) is a multifactorial disorder that affects cognitive functioning, behavior, and neuronal properties. The neuronal dysfunction is primarily responsible for cognitive decline in AD patients, with many causal factors including plaque accumulation of Aβ42. Neural hyperactivity induced by Aβ42 deposition cause abnormalities in neural networks, leading to alterations in synaptic activity and interneuron dysfunction. Even though neuroimaging techniques elucidated the underlying mechanism in the neural connectivity, precise understanding in cellular level is still elusive. Previously, a few multielectrode array studies examined the neuronal network modulation *in vitro* cultures revealing relevance of ion channels and the chemical modulators in the presence of Aβ42. In this study, we investigated neuronal connectivity and dynamic changes with high density multielectrode array, particularly in relation to network-wide parameter changes over time. By comparing the neuronal network between normal and Aβ42 treated neuronal cultures, it was possible to discover the direct pathological effect of the Aβ42 oligomer altering the network characteristics. The application of graph theory and center of activity trajectory analysis assessed the consolidation and disassociation of neural networks under Aβ42 oligomer exposure over time. This result can enhance our understanding of how neural networks are affected during AD progression.

## Introduction

Alzheimer’s disease (AD) has an adverse impact on normal cognitive functioning in memory, reasoning, and even behavior, making it the most debilitating symptom among neurodegenerative diseases[1, 2]. AD is a multifactorial disorder characterized by a multitude of factors, among which amyloid beta (Aβ) has been demonstrated to contribute significantly to its functional and morphological effects on neuronal properties [3]. In both clinical and experimental settings, paradoxical neural activity is frequently observed at different stages of AD progression [4]. The majority of patients with mild cognitive impairment (MCI) experience epileptic discharges that are followed by cognitive impairment later in life [5, 6]. Several studies suggest that pathogenic Aβ42 oligomers, formed by the endosomal proteolytic cleavage of the amyloid precursor protein (APP), cause hyperexcitability in neuronal networks [7, 8]. Induced by neural hyperactivity, Aβ42 deposition and tau spread are mutually intensifying the processes [9]. As part of the pathophysiology of Aβ42, its influence on synaptic modification has been well studied whether it causes presynaptic facilitation or postsynaptic depression [10]. Furthermore, it has been demonstrated that alterations in synaptic activity along with interneuron dysfunction may lead to network aberrations in the presence of APP or Aβ42 [11, 12]. Consequently, it is evident that Aβ42 oligomer can cause abnormalities in neural networks. To gain a more global understanding of cortical connectivity during AD progression, neuroimaging techniques such as diffusion tensor imaging, functional magnetic resonance imaging, and electroencephalography are applied clinically to dementia patients in association with their cognitive status. In contrast to the conventional method, dissociated neuron culture on HD MEA provides a more controlled means of observing neural network formation in a temporal phase, thus providing insight into neural network alteration [13]. Dissociated neuron culture on HD MEA can give insight for the neural network alteration in a more controlled manner observing network formation in spatio-temporal manner. Even though artificial, this neural culture also shares the intrinsic properties of brain, too [14]. Having acquired neural signals through HD MEA, the task of efficiently extracting topological features from big data and representing the dynamic status of the network still remains. Amongst several methodologies of connectivity analysis from brain imaging, graph theory is considered as a superior metrics informing organizational state of the neural network in terms of integration, segregation, and strength both in structural and functional levels [15-17].

In this study, the dynamic changes in neuronal networks are demonstrated under Aβ42 oligomer exposure over time. Through the analysis of a few critical network parameters using graph theory, the consolidation and disassociation of neural network are quantitatively assessed. Additionally, a center of activity trajectory (CAT) analysis is conducted to determine the distribution of network burst trajectories and the speed at which neurons transmit signals in culture arenas. It is therefore evident from the HD MEA recordings that increasing inefficiency of neural network induced by Aβ42 oligomer can facilitate a better understanding of how neural networks are affected during AD progression.

## Results

### Characteristics of cortical neuron cultures over culture ages in Aβ42 application

It is known that Aβ42 alters neuronal activity depending on its extracellular concentration, causing either presynaptic facilitation or postsynaptic depression [10]. The 10 µM Aβ42 oligomer applied in the present study has been shown to be neurotoxic in the previous study [18]. The mean firing rate (MFR) change was evident in both the visual inspection (Figure 1A) and the violin charts (Figure 1B). In both cultures, the amount of spike activity reflected from MFR was in a steady increase as it aged (Figure 1B). By comparison, MFR of Aβ42 culture was higher than control in amount and variation at least until DIV10, which implied a faster initial maturation corresponding to the previous observations of neuronal hyperactivation in Aβ42 culture [19, 20]. It is also noteworthy that the interspike interval (ISI) values as another indicator of neuronal excitability and connectivity gradually decreased in both cultures as they got older (Figure 1C). The decline pattern between Aβ42 and control cultures had a slight difference when the average ISI was dropped to 35.68% between DIV1 and 4 in control compared to 64.02% in Aβ42 culture. After DIV4, the average ISI remained relatively constant in control during which that of Aβ42 remained all time higher than control values. The difference in ISI decline pattern suggests that the Aβ42 treatment affects synaptic communication at a slower rate and lowers the receptiveness of information exchange among neurons.

**Figure 1.** Spike activation property of developing neuronal cultures on HD MEA recording measurement. (A) MFR Activity map in spontaneous neuronal activation during 5 min recordings. Control (CTL) and Aβ42 oligomer treatment groups in DIV 1, 4, 7, 10, 13 and 16. The intensity scale range from 0 to 10 spikes per second. (B) Violin plot of mean firing rate (MFR) for 5 min recordings for control (black) and Aβ42 (red) cultures during DIVs. The white dot represents the median and the thick white bar in the center represents the interquartile range. The thin black line represents the 1.5X interquartile range. (C) Box plot of interpsike interval (ISI) in millisecond for 5 min recordings for control (black) and Aβ42 (red) cultures during DIVs. The lower quartile as the borderline of the box nearest to zero expresses the 25th percentile, whereas upper quartile as the borderline of the box farthest from zero indicates the 75th percentile. Error bars show SEM. ***p□<□.005, *p□<□.05; unpaired, two-tailed t test with Welch’s correction.

### An abnormality in functional connectivity in primary cortical neuron cultures caused by Aβ42 oligomers

Figure 2A illustrates distinct dissimilarities between the control and Aβ42 cultures based on the graphical perspective of connectivity map. It is interesting to note that compared to the quantitative increase in nodal links connecting neurons in control cultures over culture ages, Aβ42 connectivity showed a biphasic pattern with an initial increase in complexity until DIV10 and then a decline until DIV16. It is similar to the neural burst and MFR observed in figure 1. CC represents the ability of neurons to form local clusters. In this respect, Aβ42 culture retained robust clustering throughout culture days with higher CC values than control (Figure 2B). Interestingly, CC values in control were consistent from DIV4 to 10 then abruptly increased from DIV13. However, PL values indicated a more dynamic change. From DIV4 to 13, the PL values in control decreased gradually (Figure 2C). Contrary to this, PL of Aβ42 culture reflected the connectivity map in a biphasic pattern with a steep incline up to DIV4 and then a gradual decline afterward. Because PL refers to the minimum number of steps necessary to reach one node from another, a shorter PL signifies more efficient connections within the network, in other words, the better small worldness [17]. In that case, the control culture developed network efficiency with age. The lowest PL at the same time as the highest CC in DIV13 of control culture implied dissociated neurons reached a state of self-organized criticality, in which a system is able to respond rapidly to changes in the environment while maintaining stability and robustness despite any external or internal perturbations [21]. However, the gradual increase of PL value from DIV4 to 10 in Aβ42 culture indicated the escalating inefficiency in network formation. The later decline in DIV13 to 16 possibly resulted from the decease of the overall network size itself noticeable both in the connectivity map as well as ND values (Figure 2A and D). As a result, Aβ42 culture did not achieve a critical state unlike control. Another network parameter, ND also provides a valuable tool to understand network organization (Figure 2D). In Aβ42 culture, due to almost 6 folds increase in average ND from DIV 4 to 7, pruning of connecting branches within nodes was not executed optimally, resulting in extra burden for neurons. NDs in control culture, however, showed a gradual increase in conjunction with a decline in PL values. According to all network parameter changes of the control culture, the network was maturing as nonessential connections were pruned, local connections were increased, and shortcuts were developed with long distance nodes. As illustrated in supplementary figure 1A and 2A, the histogram plot of the PL distribution demonstrated the maturation of neural networks in a more sophisticated manner. Firstly, Aβ42 culture demonstrated earlier neural linkage in all ranges of PL distribution compared to control cultures that initiated later. The median PL histogram for the control culture remained at 2.75 links during DIV 13 when the network reached the critical stage. On Aβ42 culture, however, the median PL histogram increased from 2.25 (DIV 7) to 3.25 (DIV 10) and 2.75 (DIV 13). The shift in median in Aβ42 culture resulted in fatter tails in the PL distribution histogram than in control, meaning the shortcut links to a distant node are longer. A noticeable growth of links in control from previous DIVs was apparent in DIV16, which kept the median value at the same level, showing the network’s integrity was stable. Aβ42 culture, on the other hand, did not display a change in the tail pattern of PL distribution, indicating that network size expansion was hampered. Similarly, the fat tail in higher ND links was also observed in Aβ42 culture (supplementary figure 1B and 2B). Based on all of the topological parameters, it is clear that Aβ42 oligomer impairs both qualitative and quantitative neural network maturation without establishing network criticality.

**Figure 2.** Neuronal connectivity map and its characterization based on graph theory. (A) neuronal connectivity map in spontaneous neuronal activation during 5 min recordings. Control (CTL) and Aβ42 oligomer treatment groups in DIV 1, 4, 7, 10, 13 and 16. Red dot represents for a node of sender, blue for receiver, and gray for broker. White line describes connection between nodes. (B) Average of clustering coefficients from neuronal connections for 5 min recordings for control (black) and Aβ42 (red) cultures during DIVs. The line represents a standard deviation. (C) Histogram plot of average path length in number of links (D) Histogram plot of node degrees in number of links.

### Modification of spike burst patterns affected by Aβ42 application

Figure 3 illustrates spike bursts in cultures that occur when neurons fire rapid action potentials within a narrow timeframe. As cultures get older, spike bursts and network bursts were observed (Figure 3A). In control cultures, bursts and frequencies were in a continuous upward trend except for DIV13 (Figure 3B and C). In contrast, spike burst duration decreased from DIV4 to DIV16 with the exception of DIV13 (Figure 3D). A similar pattern was observed for spike network bursts as well, in that their number and frequency gradually increased with a decline in duration time except for DIV13 (Figure 3E-G). The burst pattern of Aβ42 culture, however, did not show distinct characteristics. The number of spike bursts decreased from DIV1 to 4, but there was no difference between DIV4 and 7 (Figure 3B). The increase from DIV7 to 10 was reversed when it reached to DIV13. Except in DIV1, spike durations tended to decrease steadily (Figure 3D). We observed that, when DIV1 was ignored, a biphasic pattern in number and frequency of Aβ42 culture network bursts was observed that the initial incline reversed to decline at DIV10 (Figure 3E and F). In contrast, when we excluded DIV1, the network duration decreased continuously, similar to spike burst patterns (Figure 3G).

**Figure 3.** Spike burst analysis in the developing neuronal cultures. (A) Raster plot of neuronal spiking from 4096 electrodes (y axis) during 5 min recordings (x axis). Control (CTL) and Aβ42 oligomer treatment groups in DIV 1, 4, 7, 10, 13 and 16. The scale represents one minute and 400 electrodes. Mustard color represents for spike network bursts, and blue for spike bursts. (B) Box plot of number of spike burst for control (black) and Aβ42 oligomer treatment (red) groups. (C) Box plot of spike burst frequency in number of bursts per minute. (D) Box plot of spike burst duration in millisecond. (E) Line plot of number of network burst for control (black) and Aβ42 oligomer treatment (red) groups (F) Line plot of network burst frequency in number of network bursts per minute. (G) Line plot of network burst duration in millisecond. In all the box plots above, lower quartile as the borderline of the box nearest to zero expresses the 25th percentile, whereas upper quartile as the borderline of the box farthest from zero indicates the 75th percentile. Error bars show SEM. ***p□<□.005, **p□<□.01, *p□<□.05; unpaired, two-tailed t test with Welch’s correction.

### Network-wide properties of spatio-temporal signal transduction in the dissociated culture arena using CAT analysis

In the context of neuronal networks, where activity patterns are the result of complex interactions between neurons involved, the dynamics of these neurons’ firing when and where, or in other words, spatiotemporal modality, needs to be considered in a comprehensive manner. By including the vector summation of neural activities in the center of activity (CA), the flow of population activity over a short period of time can be analyzed [22]. A CA will be located closer to the center of the arena if the neural activity is more homogeneous, as the individual vector is calculated from the center of the arena. Thus, sequential tracing of CA can provide valuable information regarding the spatio-temporal trajectory of neural bursts, such as coherence, speed and dispersibility [23]. In control, there was a gradual increase in the number of network bursts as the culture aging from 0 to over 50 counts, whereas that of the Aβ42 culture is relatively constant at between 20 and 30 counts (Figure 3E). The total density of CATs was also affected by network burst count differences (Figure 4A). In control, the CA position at the end of trajectory (yellow) shifted towards the center of the arena from DIV 7 forward. A vector space can be used to visualize the travel route from beginning to end for each individual CAT (Supplementary figure 3). With the disoriented end point outside the center in Aβ42 culture, the dispersion of CAT was more spread out network-wide. Based on this result, the neural firing in control is homogeneous even at relatively early culture stages at DIV7, while the firing in Aβ42 culture remains inhomogeneous throughout the entire culture period. Aβ42 culture results in shorter CAT duration and faster velocity possibly because neural activity is localized off the center pattern of CAT (Figure 4B and C). CC values are higher in this condition as neural signals are limited to a local compartment, thereby explaining the difficulty in reaching distant neurons (Figure 2B).

**Figure 4.** CAT analysis in the developing neuronal cultures. (A) CAT analysis images of control (CTL) and Aβ42 oligomer treatment groups in DIV 1, 4, 7, 10, 13 and 16. The color scale bar represents an initiation (gray) to termination (yellow). (B) Line plot of CAT velocity with variation (vertical line) in millimeter per second (C) Line plot of CAT duration with variation (vertical line) in millisecond.

## Discussion

It is inherent for neurons to construct a neuronal network that is optimized for neuronal communications and resilience to minimize perturbations. Neurons undergo rapid axonal arborization during developmental stages, leading to random connections with other neurons through synapse formation. By removing auxiliary synapses from the preliminary network, a synaptic pruning procedure is performed to sculpt neural circuitry so that it is functional as well as energy efficient [24, 25]. Once secured, it consolidates the network structure and constantly evolves based on environmental changes, which is a key mechanism of survival in organisms. An aging brain or a neurotoxic state, which is the hallmark of Alzheimer’s disease, deteriorates network stability. Neuronal dysfunction is primarily responsible for cognitive decline in AD patients, with many causal factors including plaque accumulation of Aβ42 and tau protein. During the early stages of the disease, neuronal hyperexcitability is often seen as an indicator of the progression and severity of the disease, as it leads to the detrimental stage. As a result, understanding neuronal activity in the brain is crucial for diagnosing the condition and for studying its mechanism in an experimental setting. Previously, only a few HD MEA studies examined the relevance of ion channels and the chemical modulators of these channels in the presence of A42 [26, 27]. Here, we studied neuronal connectivity and dynamic changes, particularly in relation to network-wide parameter changes over time. By comparing the neuronal network with the same culture condition, it was possible to see the direct pathological effect of the Aβ42 oligomer. Based on CC, PL, and ND value changes, the topological and qualitative properties of the neural network were analyzed for its communication efficiency and optimization ability. In normal neuron culture, it took 13 days to build up network optimization, or, in other words, criticality. Although the quantitative network size of neurons cultured with Aβ42 oligomer was much larger than control cultures, their network became directed toward anomalies instead. Normally, the burst frequency increases while the burst duration decreased as the cultures matured, indicating a transition from a low frequency, high-duration burst pattern to a high-frequency, low-duration burst pattern. In this study, we observed the same pattern in the normal neuronal culture. On the contrary, no noticeable tendency of burst parameters could be found in Aβ42 treated culture implying the imbalance of excitatory and inhibitory synapses within the network, or changes in the intrinsic properties of the neurons. The CAT analysis also revealed a novel finding about the intrinsic nature of network structure. It had been challenging to assess the network-wide synchrony of neuronal bursts in a singular parameter containing spatiotemporal characteristics. Through CAT analysis, more homogeneous neuronal bursts were observed in the normal neurons at a relatively early stage of development. The eccentric localization of CA in Aβ42 culture indicates the fractured nature of the neural network in AD, which accounts for the scattered activation areas in other brain imaging studies.

By measuring neuronal connectivity meticulously with HD MEA, we were able to identify how neuronal connectivity developed over time. A graph theory and CAT analysis approach was employed to quantify the efficiency and synchrony of neural networks as representative parameters for defining critical states. The efficacy of therapeutic candidates can be evaluated in future studies by observing and comparing other AD factors such as tau or relevant genes on neuronal network alterations in a similar recording setting. This will allow us to screen more effective drug candidates and determine a more accurate therapeutic window to treat AD.

## Materials and Methods

### Aβ42 oligomer preparation

The peptide corresponding to human Aβ42 (Anaspec, AS-64129-1, 1 mg) was dissolved in 100 μL of DMSO by vortexing for 30 minutes at room temperature, and then the solution was added to 900 μL of PBS for incubation at 4°C for 24 hours.

### Primary neuron culture

Dissection medium Neurobasal Media (NBM) consisted of 45 ml Neurobasal Medium A, 1 ml B27 (50 X), 0.5 mM Glutamine sol, 25 μM Glutamate, 5 ml Horse serum, 500 μl penicillin/streptomycin. And culture medium consisted of 50 ml Neurobasal Medium A, 1 ml B27, 0.5 mM Glutamine sol, 500 μl penicillin/streptomycin, 50 μl HEPES. The Biochip chamber (3Brain, Arena) was cleaned, filled with 70 % ethanol for 20-30 minutes, rinsed with autoclaved DDW for 3-4 times and dried in the clean bench overnight with NBM. On the day after, 30-90 μl filtered PDLO which dissolved in borate buffer on the active surface of the Biochip was added and placed overnight in the incubator. The Biochip was washed with autoclaved DDW for 3 times before cell seeding. Primary cortical and hippocampal neuron culture were prepared from postnatal 0-day mouse pups. Pups were decapitated with sterilized scissors and the whole brain was removed. The removed brain was chilled in cold neurobasal medium with papain 0.003 g/ml solution at 4°C in a 35 mm diameter dish. Surrounding meninges and excess white matter were pulled out under the microscope (Inverted microscope, Nikon, Japan) in the same medium to a second dish at 4°C. The cortex and hippocampus parts were isolated from other parts of the brain, washed with NBM and papain solution, and minced into small pieces. The minced tissues were transferred into a 15 ml tube and incubated for 30 min in the 37°C water bath. After, the tube was inverted gently every 5 min to be mixed. The tissues were washed with HBSS twice, after being settled down, the cortex and hippocampi tissues were transferred into prewarmed NBM and triturated for 20-30 times using a fire-polished Pasteur pipette. The number of cells was counted and 30-90 µl drops of the cells were plated in the Biochip, which contains ∼1000-1500 cells/µl (incubated in 37°C in 5 % CO_2_). As a means of comparing neuronal network activity between control and Aβ42 treatment cultures over time, two sets of primary cortical neurons were prepared on 64 × 64 HD MEA chips with the same number of neurons, culture conditions, and measurement periods until DIV16. The whole medium was replaced with a fresh feeding medium every 3 days.

### Neuronal spike recording with High density Multielectrode array (HD-MEA) and data analysis

High density Multielectrode array (HD-MEA) recording with 4096 electrodes in CMOS Biochip (BiocamX, 3Brain GmbH, Switzerland) was conducted at sampling rate of 10 kHz. The active electrode which is 21 μm × 21 μm in size and 42 μm in pitch is implanted in the array with 64 × 64 grid (2.67 × 2.67 mm^2^) centered in a working area (6 × 6 mm^2^). The spontaneous neuronal spikes were recorded for 5 min at the same time of culture days. All recordings were conducted and analyzed by Brainwave software (3Brain GmbH,Switzerland). The CAT analysis in Brainwave adopted the algorithm of Gandolfo et al [23].

---

**Supplementary Figure 1**. (A) Distribution histogram of average path length in number of links to node count. Control (black) and Aβ42 oligomer treatment (red) groups in DIV 1, 4, 7, 10, 13 and 16. (B) Distribution histogram of node degree in number of links to node count.

**Supplementary Figure 2**. (A) Distribution histogram of probabilistic path length in number of links to node count percentage. Control (black) and Aβ42 oligomer treatment (red) groups in DIV 1, 4, 7, 10, 13 and 16. (B) Distribution histogram of probabilistic node degree in number of links to node count percentage.

**Supplementary Figure 3**. (A) Single representative CAT from the intermingled trajectories in figure 4. Control (CTL) and Aβ42 oligomer treatment groups in DIV 1, 4, 7, 10, 13 and 16. The color scale bar represents an initiation (gray) to termination (yellow).

## Acknowledgement

This research was supported by a grant of the Korea Health Technology R&D Project through the Korea Health Industry Development Institute (KHIDI), funded by the Ministry of Health & Welfare, Republic of Korea (grant number: HI17C1711).

