## Supplementary material for "Functional aberration of cortical neuronal network induced by Aβ42 oligomer": figure

### Slide 1
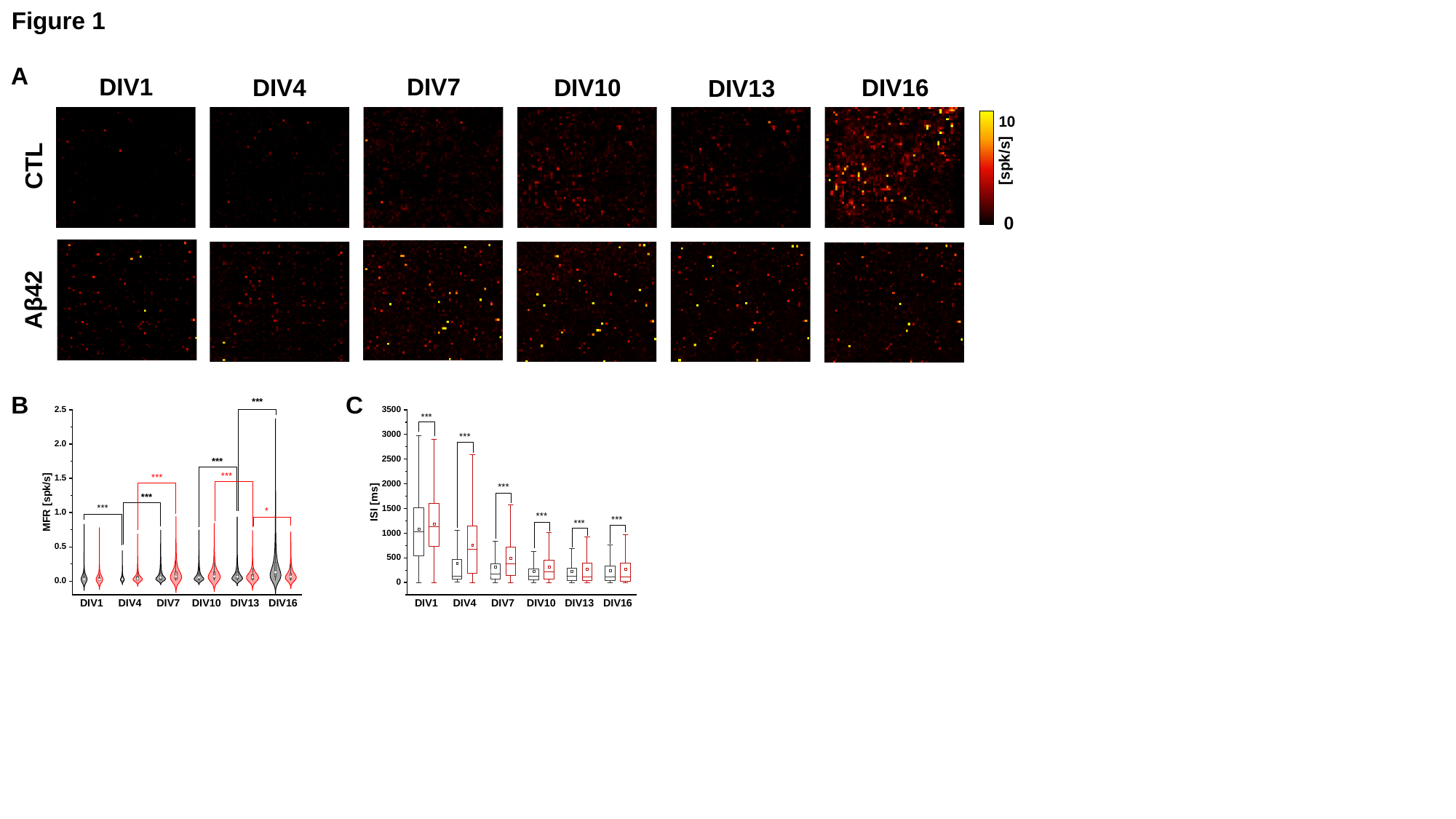

Figure 1
A
DIV1
DIV7
DIV4
DIV10
DIV16
DIV13
10
[spk/s]
CTL
0
Aβ42
B
C

### Slide 2
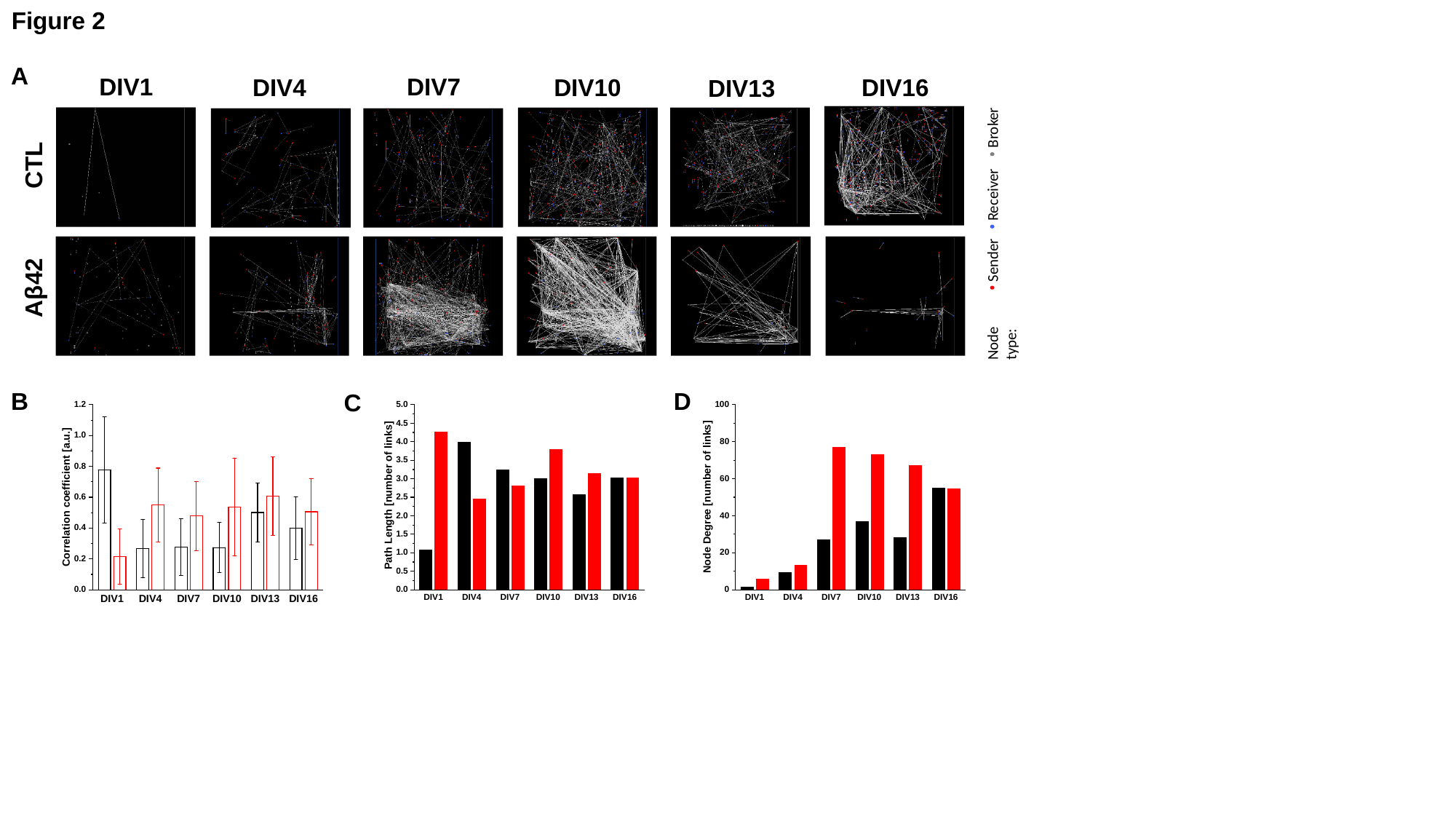

Figure 2
A
DIV1
DIV7
DIV4
DIV10
DIV16
DIV13
CTL
Sender
Receiver
Broker
Node type:
Aβ42
B
D
C

### Slide 3
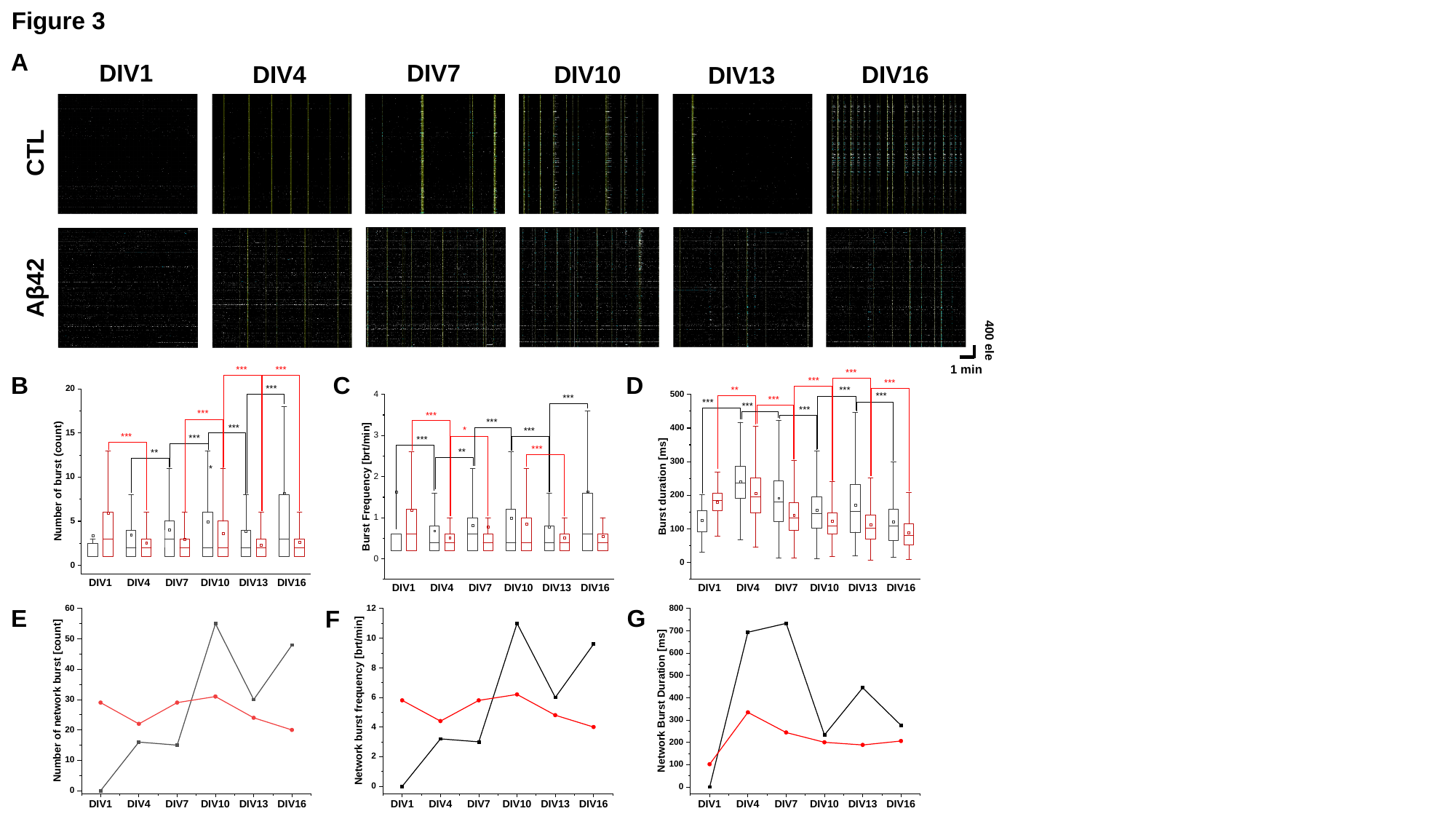

Figure 3
A
DIV1
DIV7
DIV4
DIV10
DIV16
DIV13
CTL
Aβ42
400 ele
1 min
B
C
D
E
G
F

### Slide 4
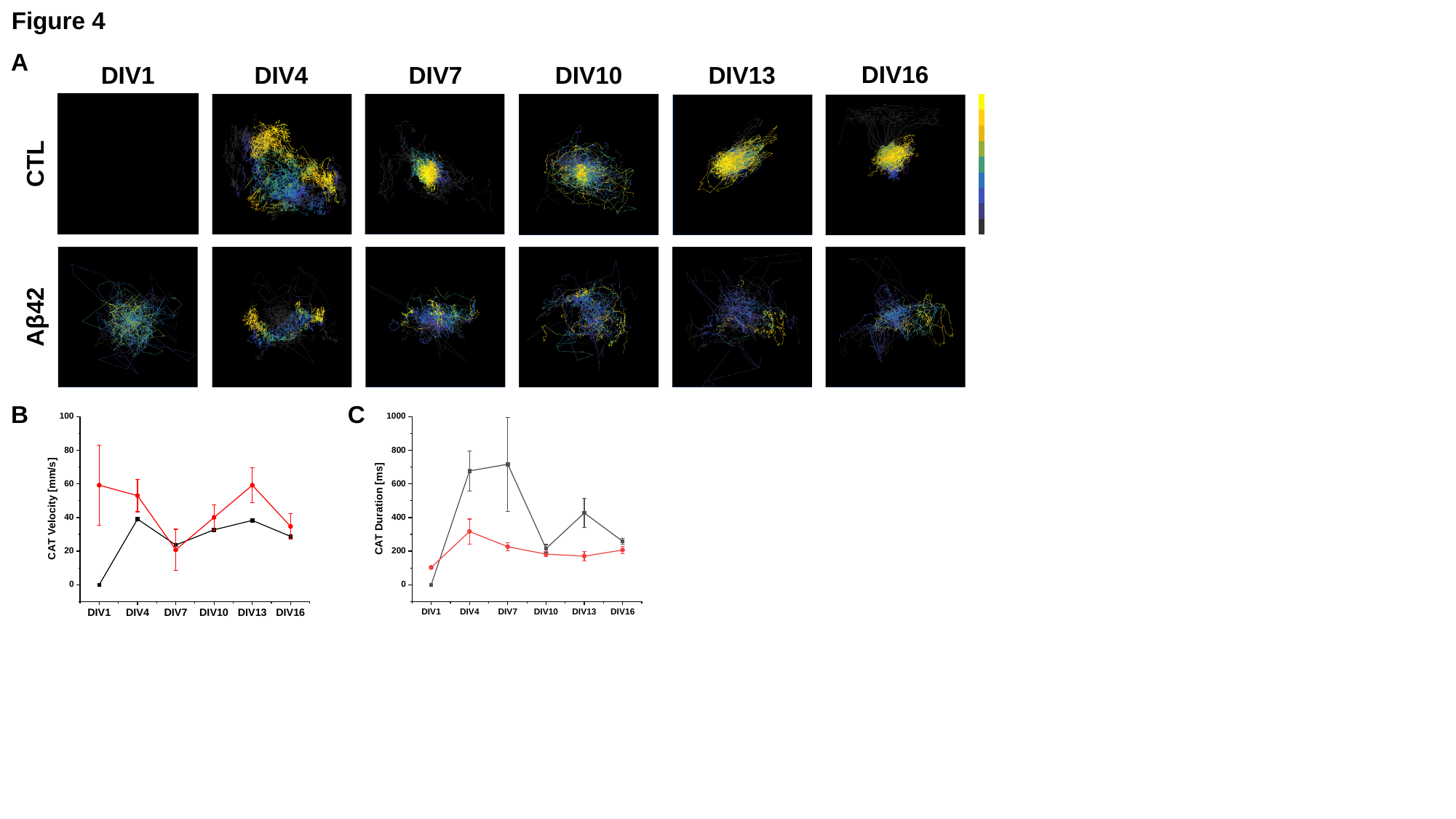

Figure 4
A
DIV16
DIV1
DIV4
DIV7
DIV10
DIV13
CTL
Aβ42
B
C

### Slide 5
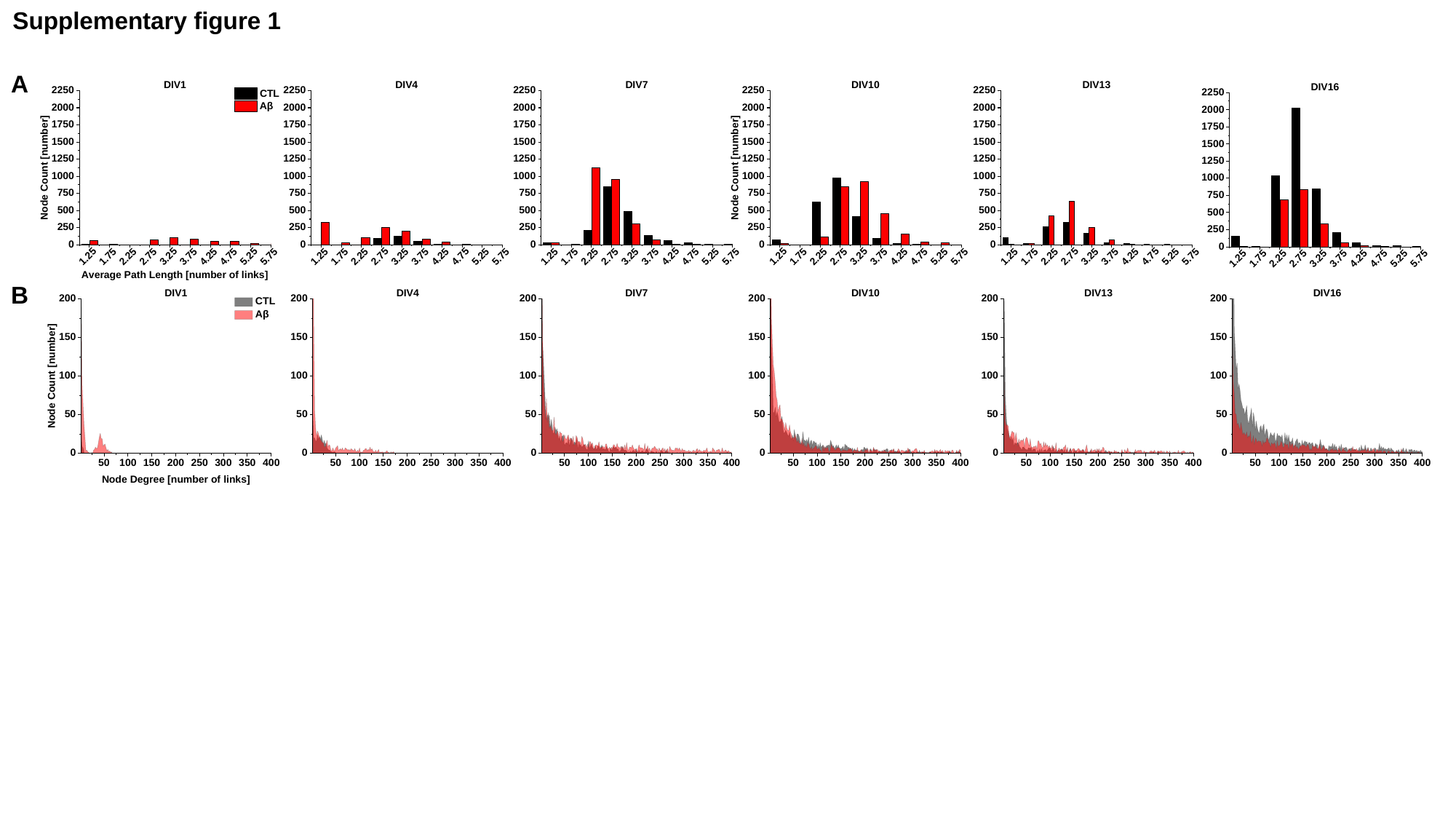

Supplementary figure 1
A
B

### Slide 6
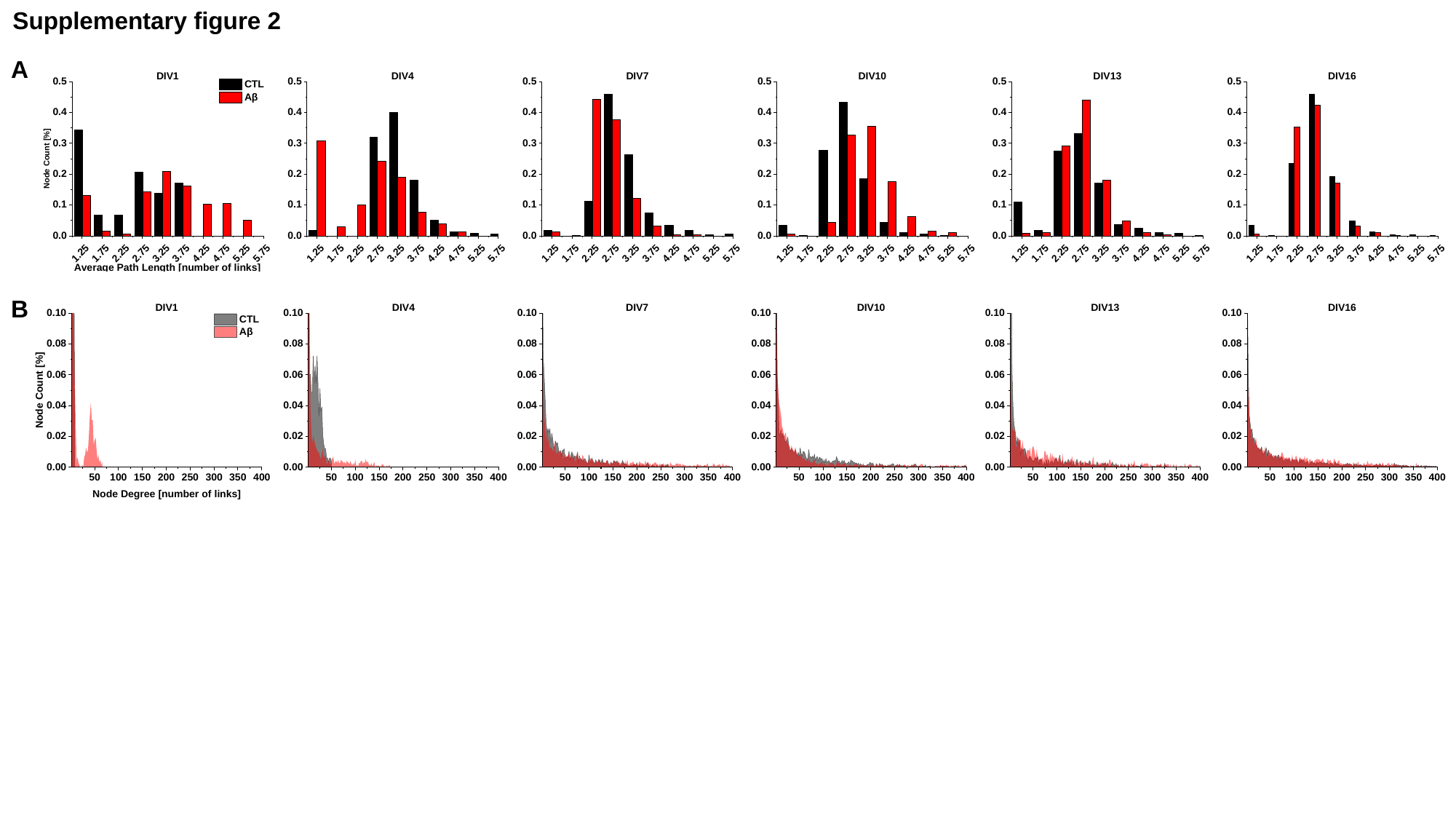

Supplementary figure 2
A
B

### Slide 7
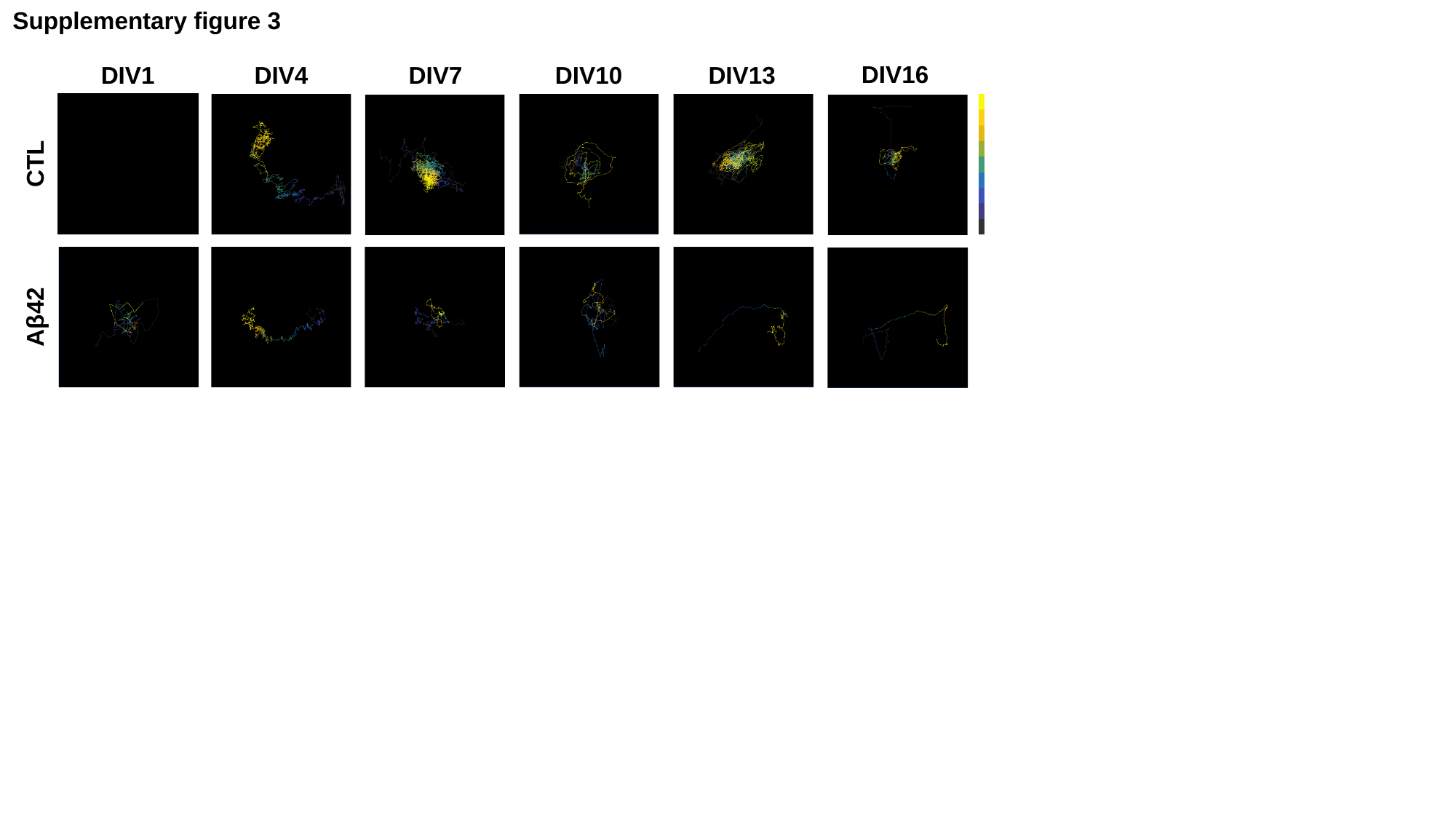

Supplementary figure 3
DIV16
DIV1
DIV4
DIV7
DIV10
DIV13
CTL
Aβ42
